# How developmental constraints shape the evolution of repeated structures

**DOI:** 10.1101/2025.03.27.645690

**Authors:** Daohan Jiang, Matt Pennell, Lauren Sallan

## Abstract

Repeated structures are widespread across multicellular organisms, such as vertebrae, segments, and cell types. These structures, also known as serial homologs, share ancestral states and developmental underpinnings yet also provide the materials for novel adaptive phenotypes. It remains largely unclear why some repeated structures diverge quickly, while others remain constrained. One reason for this uncertainty is the lack of a generalized model for the evolution of repeated structures under different scenarios that links developmental, genetic, and selective constraints to expected and observed patterns of evolution. Here, we introduce a model that incorporates key structural features of gene regulatory networks and selection and investigate how responses to multivariate selection depend on developmental constraints. We show structural features of developmental networks determine when repeats can respond independently to selection and when divergence is limited. Simulations recover broad expectations of phenotypic evolution under directional selection inferred from empirical data. We further show that, in the face of fluctuating selection, strong developmental constraints lead to reduced fitness over time and attenuated fitness fluctuations. Together, our results provide general insights into the principles of evolution of repeated structures and offer a modeling framework for the evolution of a broad range of key phenotypic characters.

## Introduction

The body plans of all multicellular organisms feature a number of repeated structures with recognizable phenotypic similarity and shared ancestral states, whether in adults, during early stages of development, or over evolutionary history. The divergence of repeated structures (also known as serial homologs; "repeats" hereafter) within an organism is an essential means by which novel, adaptive phenotypes have evolved [1–3]. Notable examples of repeats parts that develop into distinct forms include paired fins and limbs of vertebrates, vertebrae, gill arches, skin appendages (e.g., scales, feathers, hair), body segments of arthropods and paired appendages attached to the segments, components of angiosperm flowers, and cell types [3].

Central to the problem of the evolution of repeats is how developmental processes can constrain and bias the trajectory of evolution. Developmental constraints can have a dispositional effect on phenotypic evolution as developmental processes channel the effect of mutations on the phenotype and bias the availability of phenotypic variation [4–13]. Developmental constraints are particularly relevant to phenotypic divergence between repeats, as they necessarily share underlying developmental processes at the point of origination, introducing constraints that must be broken to enable modification of any particular copy. While homologous structures of different species (e.g., forelimbs of different tetrapod species) can undergo phenotypic divergence as different lineages acquire different genomic substitutions, repeats within the same organism (e.g., forelimbs and hindlimbs of the same animal) can only take distinct forms if the originally identical developmental processes become different [3, 14–20]. The evolutionary divergence of repeats, therefore, needs to be understood in the light of development.

There has, however, been little systematic study of how and when repeats diverge. Our understanding of the principles of their developmental evolution has been limited by the absence of relevant, broad empirical data which can set baseline expectations, namely full phenotypic transformation series for different parts with known molecular underpinnings and developmental origins. Such data, obtained from fossils, embryos, and outgroups, can help set baseline expectations for the rate and mode of evolution [21–25]. In the absence of such datasets, an alternative approach is to build frameworks of evolutionary simulations based on core mechanistic assumptions about developmental evolution, in order to test their plausibility and understand how phenotypic evolution might proceed under different scenarios. Such a modeling framework, when combined with relevant data, can also help us infer possible regimes of selection that best account for the observations. This approach has been applied to specific morphological characters with well-understood developmental biases, such as mammalian molars, allowing improved inference of divergence patterns and rates (e.g., [26]). However, such detailed understanding of developmental mechanisms is usually unavailable for phenotypic traits or taxa of interest, and generalizable models are necessary before sufficient details of development are revealed.

Here, we have constructed an evolutionary model that incorporates general features of gene regulatory networks (GRNs) in order to establish and test basic assumptions about the frequency and speed of phenotypic divergence in repeats under different scenarios of selection and developmental constraint. Under this model, the phenotypic state of each repeat is controlled via a hierarchical gene regulatory network consisting of two classes of genes: the regulatory genes and the effector genes. The effector genes are responsible for producing the phenotype, and a common set of effector genes is expressed in all repeats, at least ancestrally. The regulatory genes, on the other hand, encode *trans*-regulatory factors like transcription factors (TFs) that interact with *cis*-elements (e.g., promoters and enhancers) of the effector genes and regulate the expression of the effector genes [3]. Importantly, as the same effector genes are regulated by different *trans*-factors in different repeats during development, mutations in the *cis*-elements can potentially have different local effects on effector gene expression in different repeats. If regulation of effector gene expression relies solely on pleiotropic *cis*-elements that interact with *trans*-factors expressed in multiple repeats, and the *trans*-factors have similar binding preferences (e.g., paralogs with very similar sequences), *cis*-regulatory mutations are likely to have concordant effects on different repeats, resulting in genetic correlation between their phenotypic states. In contrast, if the *trans*-factors have dissimilar binding preferences, or if there are non-pleiotropic, organ-specific *cis*-elements, the effects of *cis*-regulatory mutations on different repeats would be more decoupled. In this study, we focus on evolutionary changes in the *cis*-elements and their interactions with the TFs, as evolutionary changes in protein sequences or expression levels of the TFs tend to have greater pleiotropic effects and are generally less likely to contribute to phenotypic evolution and adaptation [27, 28].

Under this model, we derived the expected coevolutionary dynamics under multivariate selection on the phenotype, and examined how the sharing of regulatory apparatus between repeats could constrain the dynamics of phenotypic evolution and phenotypic response to directional and fluctuating selection, demonstrating that the extent of coupling between gene regulation in different body parts can result in differences in the rate of adaptation and fitness over the long term in a changing environment.

## Results

### Response to directional selection under developmental constraints

We modeled the evolution of two repeated organs, with the development of each repeat regulated by a unique TF. Phenotypic states of the two repeats (denoted as *z*_1_ and *z*_2_, respectively) are produced by the same set of effector genes, whose expression during development is regulated via the interaction between the TFs and their *cis*-elements. A *cis*-element could be either pleiotropic (interacting with TFs in both repeats) or non-pleiotropic (accessible to a TF in only one of the repeats but not in the other). A mutation in non-pleiotropic *cis*-element affects gene regulation in only one body part, resulting differential expression of effector genes and differences in adult phenotypes between body parts (Fig. 1).

**Figure 1:**
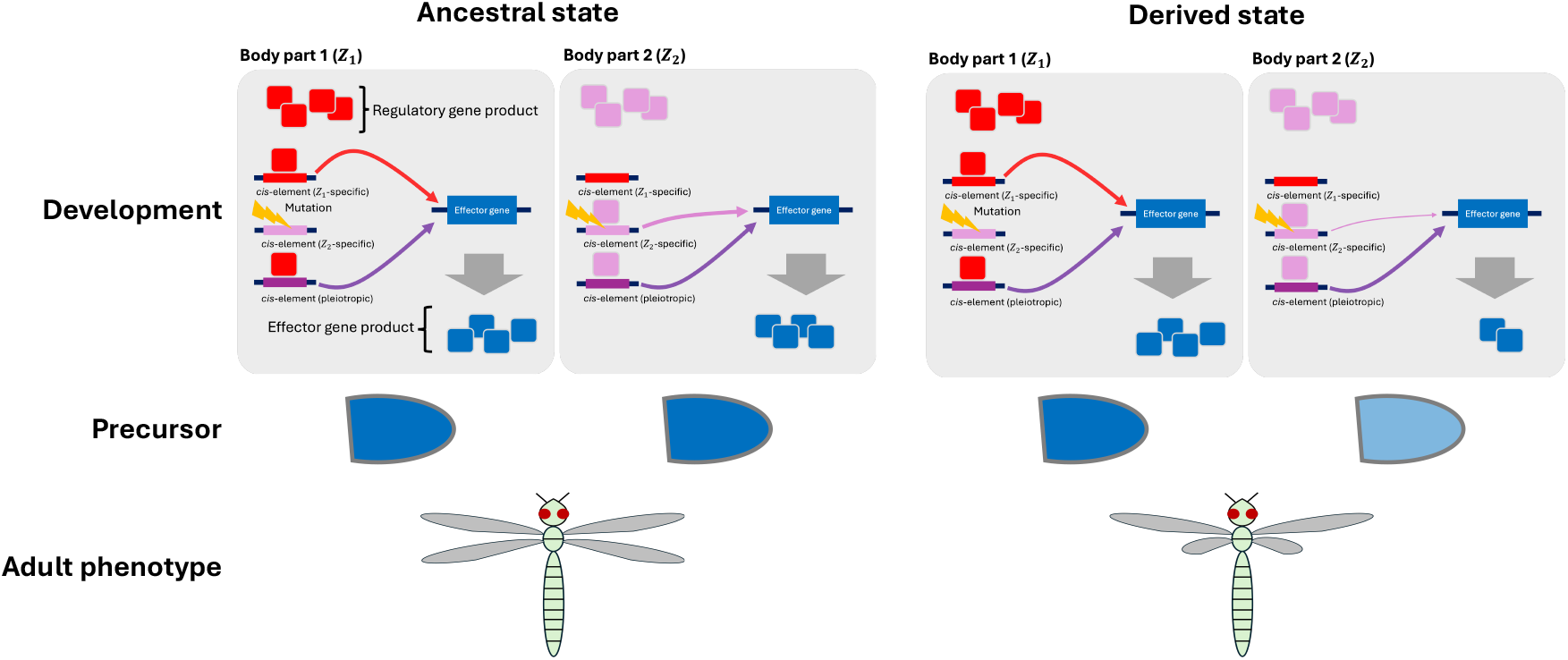
Schematic illustration of evolutionary divergence between two repeated body parts (*Z*_1_ and *Z*_2_), where two distinct regulatory genes are expressed during development. Transcription factors (TFs) encoded by the two regulatory genes activate the same effector gene(s) via *Z*_1_-specific, *Z*_2_-specific, or pleiotropic *cis*-elements. Expression of the effector gene(s) determines the phenotypic state (i.e., length of wings). A *cis*-mutation in a *Z*_2_-specific *cis*-element is shown; the mutation only affects *cis*-*trans* interactions in *Z*_2_ and only the phenotypic state of *Z*_2_. Upper row: developmental regulation in two body parts. Middle row: the developmental precursor (outgrowth) that would grow into , with color corresponding to concentration of the effector gene product. Lower row: character state in the adult organism. The ancestral state is shown on the left side, and the derived state with the depicted mutation fixed is shown on the right side.

Under this model, we derived how developmental parameters shape the mutational variance-covariance matrix (the **M**-matrix), which can be verified with mutation accumulation (MA) simulations (Fig. S1). With the **M**-matrix, the evolution of the population mean phenotype under the joint effect of mutation, genetic drift, and selection can then be modeled as a multivariate Ornstein–Uhlenbeck (OU) process. Using simulations, we explored the dynamics of evolution under three scenarios of directional selection, in all of which a greater value of *z*_1_ is selected for. In the first scenario, *z*_2_ is under stabilizing selection (Fig. 2A, left); the response of *z*_1_ to directional selection reflects the conditional evolvability [29]. In the second scenario, there is selection on the proportion between *z*_1_ and *z*_2_ (i.e., stabilizing selection on *z*_1_ *− z*_2_) and greater values of both *z*_1_ and *z*_2_ are selected for (Fig. 2A, middle); in this scenario, the selective line of least resistance (SLLR; the principal axis of the adaptive landscape and the phenotypic dimension where selection is the weakest [30, 31]) is aligned with the genetic line of least resistance (GLLR; the phenotypic dimension where genetic variance is produced most quickly [32]). In the last scenario, there is a functional tradeoff between *z*_1_ and *z*_2_ (i.e., stabilizing selection on *z*_1_ + *z*_2_) and a smaller value of *z*_2_ is selected for; in this scenario, SLLR is orthogonal to GLLR (Fig. 2A, right). In the latter two scenarios, *z*_1_ and *z*_2_ are under correlational selection. We specifically examined how the proportion of *cis*-elements that are pleiotropic (*f*_*p*_) and the correlation between pleiotropic mutations’ effects on two body parts (*ρ*) affect the evolutionary dynamics of *z*_1_. We show that, given a high *ρ*, response of *z*_1_ to selection becomes slower as the proportion of *cis*-elements that are pleiotropic (*f*_*p*_) increases in the cases where there is stabilizing selection on *z*_2_ (Fig. 2B, left) or a functional tradeoff between body parts (Fig. 2B, right). In contrast, when selection on proportion is present, *f*_*p*_ has a positive effect on adaptation (Fig. 2B, middle). When binding preferences of the TFs are only moderately similar such that pleiotropic mutations’ effects on *z*_1_ and *z*_2_ are less strongly correlated, the effect of *f*_*p*_ becomes less pronounced, albeit qualitatively similar (Fig. S2).

**Figure 2:**
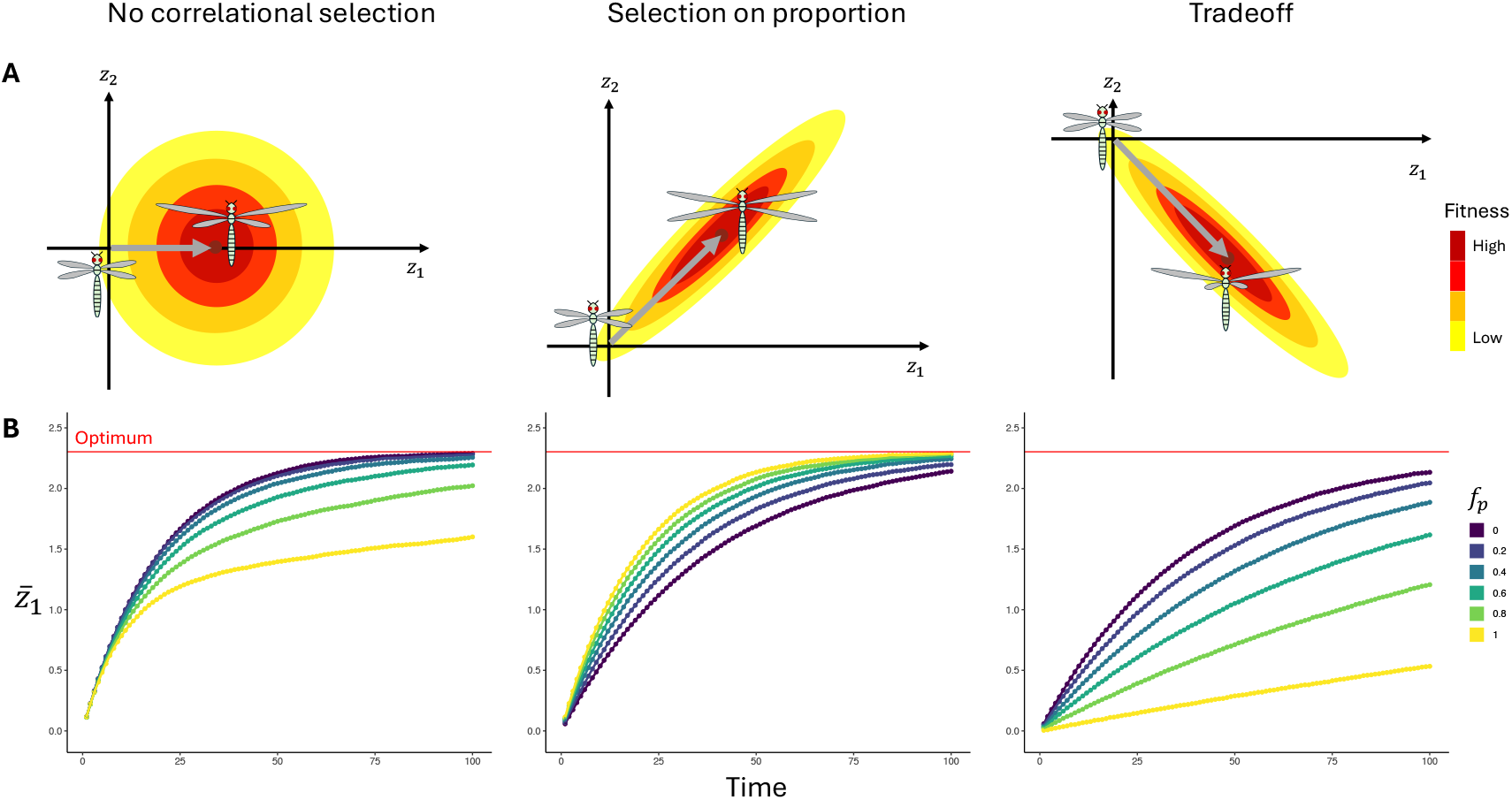
Response of the phenotype to directional selection. (A) Schematic illustration of three regimes of selection. The axes correspond to phenotypic states of two body parts (*z*_1_ and *z*_2_), and the origin point is the ancestral state. The ellipses in each panel together represent the adaptive landscape near the optimal phenotype. The optimum (adaptive peak) is at the center of the ellipses and fitness declines with the distance to the optimum, which is denoted as the color gradient from red to yellow. The gray arrow in each panel starts at the ancestral state and points to the optimum, representing the overall direction of selection. In all scenarios, a value of *z*_1_ that is greater than the ancestral value is selected for. Left: only *z*_1_ is under directional selection while *z*_2_ is under stabilizing selection. Middle: there is selection on the proportion between *z*_1_ and *z*_2_ (*z*_1_ = *z*_2_ is the optimal proportion) and a greater value of *z*_2_ is selected for. Right: there is a functional tradeoff between *z*_1_ and *z*_2_ (*z*_1_ + *z*_2_ = 0 is the optimal allometric relationship), and a smaller value of *z*_2_ is selected for. (B) Mean of *z*_1_ across simulated populations (denoted as 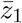) over time (steps), with the left, middle, and right panels corresponding to scenarios of selection in the left, middle, and right panels in (A), respectively. Colors of the curves correspond to scenarios with different proportions of *cis*-elements that are pleiotropic (*f*_*p*_). The results shown here were obtained with *N*_*e*_ = 10^5^ and high similarity in binding preference between TFs expressed in two body parts (correlation coefficient between pleiotropic mutations’ effects on *z*_1_ and *z*_2_ is *ρ* = 0.9; see Methods).

The above results rely on the assumption that selection is weak and the effective population size is reasonably large [33]. To see how general the effect of *f*_*p*_ is in constraining adaptation, we performed individual-based population genetic simulations where these assumptions do not hold. The effect of *f*_*p*_ on adaptation is qualitatively similar (Fig. S3), indicating generality of the effect of developmental constraint on evolution.

### Delayed tracking of adaptive peak movement under developmental constraints

Next, we examined how developmental constraints could influence responses to fluctuating selection as imposed by a changing environment. Specifically, we asked how *f*_*p*_ would affect the population’s ability to track an adaptive peak (optimal phenotype) as it moves over time. We first considered scenarios where the peak moves continuously like a Brownian motion (BM) process, which is expected to result in BM-like evolutionary dynamics given that the populations should closely track the adaptive peak [34]. We considered the same three regimes of selection on the phenotype as depicted in Fig. 2A, but let the peak move over time. It is assumed that the movement of the peak is constrained by internal selection such that the main axis of peak movement is aligned with SLLR [31]: in the absence of correlational selection, optima of *z*_1_ and those of *z*_2_ over time will be uncorrelated (Fig. 3A, left panel); when correlational selection is present, the main axis of movement is aligned with SLLR (Fig. 3A, middle and right panels). As a result, the direction of peak movement is generally aligned with GLLR when there is selection for proportion between *z*_1_ and *z*_2_, but orthogonal to GLLR when there is a functional tradeoff between them.

**Figure 3:**
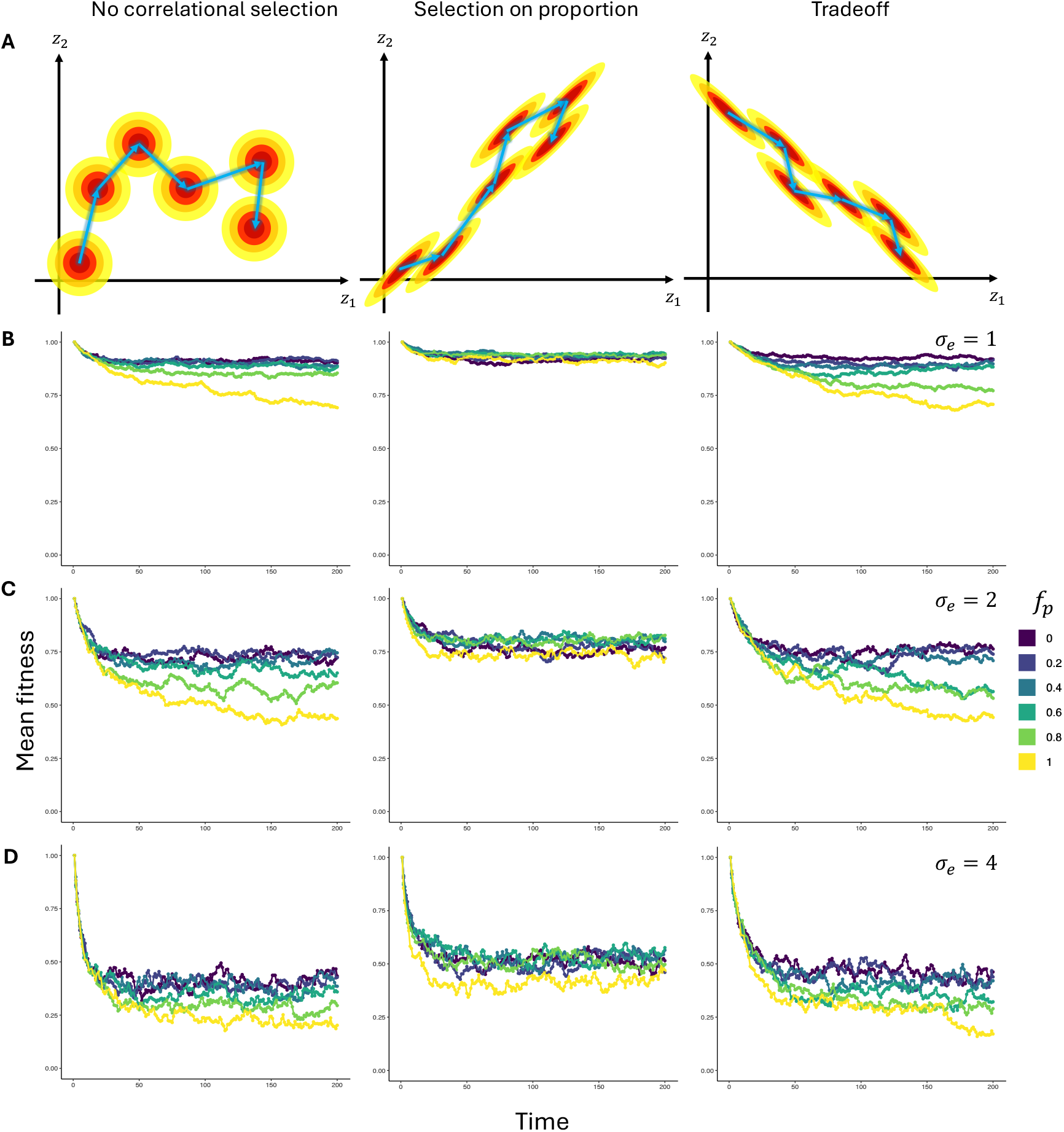
Evolutionary dynamics under fluctuating selection, where the adaptive peak (optimal phenotype) shifts over time according to a bivariate Brownian motion (BM) model. (A) Schematic illustrations of regimes of adaptive peak movement. Each panel shows multiple episodes of peak movement in a lineage and the adaptive landscape near each peak. Each arrow between peaks represents the direction of the shift. Left: no correlational selection is present, and movements of the optimal values of *z*_1_ and *z*_2_ are uncorrelated. Middle: there is selection on the proportion between *z*_1_ and *z*_2_, and the adaptive peak generally moves along the *z*_1_ = *z*_2_ line. Right: there is a functional tradeoff between *z*_1_ and *z*_2_, and the adaptive peak generally moves along the *z*_1_ + *z*_2_ = 0 line. (B-D) Mean fitness of simulated populations over time. The left, middle, and right panels correspond to regimes of adaptive peak movement shown in the left, middle, and right panels of (A), respectively. (B) The adaptive peak moves at a relatively slow rate (*σ*_*e*_ = 1). (C) The adaptive peak moves at an intermediate rate (*σ*_*e*_ = 2). (D) The adaptive peak moves at a relatively fast rate (*σ*_*e*_ = 4).

For each combination of *f*_*p*_, rate of peak movement (*σ*_*e*_), and mode of selection (and thereby peak movement), we simulated 100 replicate lineages, whose adaptive peaks moved independently over time. The overall adaptive performance of the replicate lineages at a given time point was calculated by first calculating the fitness of each lineage as a function of their respective mean and optimal phenotypes at the time and then calculating the average fitness across lineages. Simulations show that, in scenarios where the main axis of peak movement was not aligned with GLLR, lineages with high *f*_*p*_ generally had lower fitness over time (Fig. 3B-D, left and right panels), and geometric mean of fitness over time is negatively correlated with *f*_*p*_ (Fig. S4, left and right panels). In scenarios where the adaptive peak moved along SLLR, the effect of *f*_*p*_ is minimal (Fig. 3B-D, middle panels) and the correlation between geometric mean fitness and *f*_*p*_ is not significant (Fig. S4, middle panels). Nonetheless, geometric mean fitness varied considerably among lineages with the same *f*_*p*_ in all scenarios (Fig. S4), indicating the performance of a lineage is contingent on the specific trajectory of movement of the adaptive peak.

Despite inability of the lineages to track the adaptive peak in many scenarios, phenotypic variance across lineages consistently increased over time, with the correlation coefficient between *z*_1_ or *z*_2_ and time above 0.95 in all scenarios (Fig. S5). Hence, BM-like evolutionary dynamics and a high phylogenetic signal can result from not only neutral evolution or close tracking of the adaptive peak, but also largely imperfect, lagged tracking. Correlation between *z*_1_ and *z*_2_ across lineages, on the other hand, gradually approached the correlation between movements of their optima, thereby deviating from the genetic correlation when GLLR and the main axis of optimum movement are poorly aligned (Fig. S6), indicating that developmental constraints can influence the alignment between genetic correlation and phenotypic evolution. Together, we show that the presence of developmental constraints can potentially limit adaptation in a changing environment and leave a predictable signature on the dynamics of trait coevolution.

### Complex impact of developmental constraints in the face of abrupt adaptive peak shifts

We considered an alternative regime of fluctuating selection, which we refer to as a white noise (WN) model: a new adaptive peak is resampled from a pre-specified multivariate distribution periodically; under this model, the new optimum is independent of the previous one (Fig. 4A). Again, we considered the same three regimes of selection on the phenotype as depicted in Fig. 2A, and assumed the movement of the adaptive peak to be constrained by internal selection: when there is selection for proportion, the distribution from which new optima are sampled will have a positive covariance and the main axis of peak movement will be aligned with GLLR; when there is a tradeoff between *z*_1_ and *z*_2_, the distribution will have a negative covariance and the main axis of peak movement will be orthogonal to GLLR.

**Figure 4:**
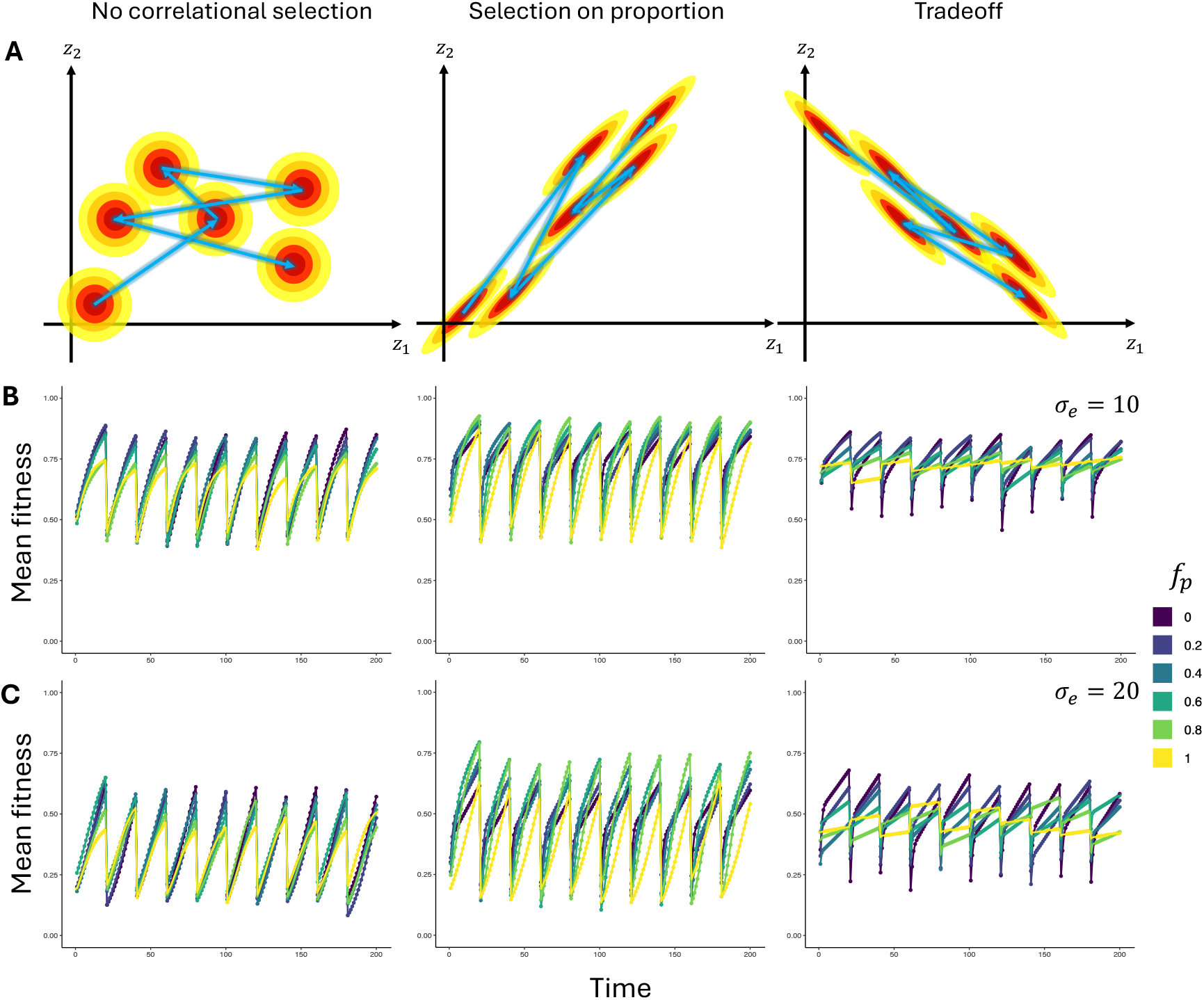
Evolutionary dynamics under fluctuating selection, where the adaptive peak shifts over time in a memoryless, white noise (WN)-like fashion. The optimum is resampled periodically from a pre-specified bivariate normal distribution. (A) Schematic illustrations of regimes of adaptive peak movement. Each panel shows multiple episodes of peak movement in a lineage and the adaptive landscape near each peak. Each arrow between peaks represents the direction of the shift. Left: no correlational selection is present, and changes in the optimal values of *z*_1_ and *z*_2_ upon each shift are uncorrelated. Middle: there is selection on the proportion between *z*_1_ and *z*_2_, and the main axis of adaptive peak movement is the *z*_1_ = *z*_2_ line. Right: there is a functional tradeoff between *z*_1_ and *z*_2_, and the main axis of adaptive peak movement is the *z*_1_ + *z*_2_ = 0 line. (B-C) Mean fitness of simulated populations over time. The left, middle, and right panels correspond to regimes of fluctuating selection in the left, middle, and right panels of (A), respectively. (B) The optimum is resampled from a relatively narrow distribution (standard deviation *σ*_*e*_ = 10). (C) The optimum is resampled from a relatively wide distribution (standard deviation *σ*_*e*_ = 20).

Unlike in scenarios where peak movement is more continuous, fitness fluctuated periodically, reflecting episodes of response to directional selection (Fig. 4B-C). When there is no correlational selection, lineages with high *f*_*p*_ generally showed slower response to directional selection and had lower fitness at the end of each episode of adaptation (Fig. 4B-C, left panels). When there is selection for proportion, interestingly, it is intermediate *f*_*p*_ that led to the highest fitness after each episode of adaptation (Fig. 4B-C, middle panels). When there is a tradeoff between *z*_1_ and *z*_2_, high *f*_*p*_ notably resulted in fitness fluctuation at a smaller magnitude, whereas lower *f*_*p*_ resulted in both high fitness at the end of each episode of adaptation and low fitness immediately after each shift of the optimum (Fig. 4B-C, right panels). Despite the apparent stability of fitness over time associated with high *f*_*p*_, a weak but significant negative correlation between *f*_*p*_ and geometric mean fitness and considerable variation among lineages were observed in all scenarios examined (Fig. S7). Correlation between *z*_1_ and *z*_2_ across lineages generally approached the correlation between their optima but also showed periodic fluctuation (Fig. S8).

## Discussion

In this study, we present a generalizable model of developmental evolution and use it to explore the evolutionary dynamics of repeated organs. We focused on a setting where different body parts had a shared set of effector genes but distinct regulatory genes expressed during development, with phenotypic divergence between body parts mediated by *cis*-regulatory substitutions that have differential effects on effector expression in different body parts. This type of evolutionary change is relatively common and contributes more to phenotypic variation and evolution compared to changes at other layers of the developmental hierarchy [27, 28]. Under this setting, we show that the coupling of gene regulation in different body parts can have a marked effect on the evolutionary dynamics. When *f*_*p*_ is high, which means the expression of the effector genes is mainly regulated via pleiotropic *cis*-elements, the resulting genetic correlation between body parts can constrain adaptation if the direction of selection is misaligned with the GLLR (Fig. 2, left and right panels). In contrast, when there is selection for proportion between body parts such that the direction of selection is aligned with GLLR, pleiotropy of *cis*-elements instead facilitates adaptation (Fig. 2, middle panels).

To investigate how the constraint resulting from the pleiotropy of *cis*-elements could affect evolvability in a changing environment, we examined the evolutionary dynamics under two regimes of fluctuating selection where the adaptive peak shifts over time. Under the first regime of fluctuating selection, where the adaptive peak moves continuously over time in a BM-like fashion, high *f*_*p*_ results in lower fitness over time if the main axis of peak movement is misaligned with GLLR (Fig. 3, left and right panels); when the main axis of peak movement is aligned with GLLR, however, *f*_*p*_ does not have a positive effect on fitness over time either. In the other regime of fluctuating selection, where the adaptive peak shifts abruptly in a WN-like fashion, *f*_*p*_ displays more complex interactions with the mode of correlational selection. In the absence of correlational selection, high *f*_*p*_ results in lower fitness at the end of each episode of adaptation. When the main axis of peak shift is aligned with GLLR, an intermediate *f*_*p*_ maximizes fitness at the end of each episode. At last, when the main axis of peak movement is orthogonal to GLLR, high *f*_*p*_ results in less fluctuation of fitness over time.

In this study, we considered relatively simple settings where the regulatory genes themselves, their expression, and the number of *cis*-elements that can potentially channel their regulatory effects, all remain constant over time. Evolutionary changes in these aspects of the GRN, while relatively rare compared to substitutions in the *cis*-elements, can have a profound effect on the structure of phenotypic variation and thereby long-term evolvability [3, 18, 28, 35, 36]. The emergence of new organ or tissue-specific *cis*-elements, for instance, can reduce *f*_*p*_ and thus decouple evolutionary changes in different body parts. It is of great interest to future studies how selection for evolvavility, especially that in a changing environment, would shape the coevolutionary dynamics of different repeats, though parameterization of the mutational origin of novel *cis*-elements can be challenging, and would require more knowledge about the regulatory potential of putative *cis*-elements and selection on these sequences (e.g., [37–42]. Divergence in binding preferences between the TFs expressed in different repeats can also decouple evolutionary changes in the repeats by making *f*_*p*_ less consequential. However, as regulatory genes are often pleiotropic and involved in the development of multiple organs, mutations that alter their binding preferences are likely strongly deleterious [27], unless the effect is localized (e.g., [43]).

Another type of evolutionary change that is not considered in the present study, but of potential interest to future studies, is the evolution of novel regulatory gene expression programs. These are hypothesized as playing a key role in evolutionary innovations, including in the origin of repeats with distinct identities in the first place [3]. As each regulatory gene can regulate a number of effector genes, evolutionary change in their expression can have drastic effects on the phenotype. Furthermore, the expression of additional regulatory gene(s) can also result in a change in variability and unlock a new phenotypic subspace by causing a set of otherwise silent effector genes to be expressed in a body part or tissue. The ectopic expression of such effector genes could make an otherwise invariable phenotypic dimension variable; such a phenotypic dimension is an evolutionary novelty that is unique to the focal lineage and the focal body part. Such novelties can complicate the application of the model presented in this study to empirical data. The model is readily applicable if the body parts of interest share a recognizable structural "plan" such that "corresponding" phenotypic dimensions that are likely controlled by the same effector genes can be identified; in contrast, phenotypic dimensions that were not present in both body parts ancestrally cannot be represented by traits in the simplistic model used in the present study. For instance, forewings and hindwings of damselflies have a shared venation pattern [44, 45], and it can be assumed that morphometric landmarks placed at matching locations are affected by the same effector genes, regulated by TFs specifying forewing and hindwing identities in forewing and hindwing development, respectively [16, 46–53]. In contrast, the forewings and hindwings of dragonflies have different venation patterns, presumably reflecting the involvement of regulatory genes specifying evolutionarily novel wing areas [44]. In some more extreme cases, one or more of the repeats have become so different from the ancestral form that "corresponding" phenotypic dimensions are no longer identifiable (e.g., elytra of beetles and halteres of flies), the model used in this study would not be applicable to modeling their phenotypic coevolution. Furthermore, our model applies to repeats with the same ancestral developmental program and phenotype, but not body parts that acquired phenotypic similarity due to secondary co-option. For instance, eye spots on butterfly wings share developmental TFs with the legs because the same TF was co-opted [54], in which case our model is not applicable.

Together, we present a generalizable model for the evolution of repeated structures of multicellular organisms, incorporating their underlying developmental mechanisms. Using this model, we demonstrate the interplay of selective and developmental constraints in shaping the evolutionary dynamics. Readily ex-tendable to a broader range of morphological characters, our model offers a framework for understanding the principles of phenotypic evolution and evolvability.

## Methods

### Developmental constraints on phenotypic variability

Let us consider two repeated organs (referred to as *Z*_1_ and *Z*_2_ respectively). Their identities are distinguished by the expression of regulatory genes (TFs) *R*_1_ and *R*_2_, respectively, during morphogenesis. For simplicity, regulatory genes that are expressed in both repeats are not considered for now and will be omitted in the formulations below. The omitted regulatory genes do not contribute to phenotypic differences between the focal repeats but may contribute to that between the focal repeats and other body parts.

Let us consider a phenotypic trait whose value results from post-morphogenesis growth. The phenotypic effect of each regulatory gene on the trait value is mediated by a set of *cis*-element, which the regulatory gene interacts with during development. Let there be *n cis*-elements that can potentially interact with *trans*-factors and contribute to the phenotype. The phenotypic state of *Z*_1_, denoted as *z*_1_, is given by 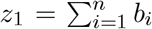 ln *a*_1,*i*_, where *a*_1_ *>* 0 is the strength of interaction between the gene product of *R*_1_ and the *i*-th *cis*-element, and *b*_*i*_ represents the phenotypic effect of the *cis*-element on the phenotype. Similarly, for *Z*_2_, there is 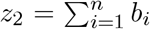 ln *a*_2,*i*_. When *b*_*i*_ *>* 0, an interaction between the *i*-th *cis*-element and the TF has a positive effect on the trait value; when *b*_*i*_ *<* 0, the effect is negative. Biologically, *b*_*i*_ encompasses the regulatory effect of the *cis*-element on the downstream effector genes (e.g., if it is a promoter, enhancer, or silencer) and the effect of the effector genes on the phenotype (e.g., whether they facilitate or repress growth).

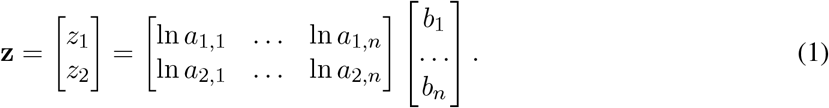

Note that *z*_1_ and *z*_2_ represent normalized phenotypes (e.g., log fold difference from a reference value), whereas their exponential (e.g., exp *z*_1_ and exp *z*_2_) can represent phenotypic traits that take only positive values, such as length or mass. When 0 *< b*_*i*_ *<* 1 and parameters other than *a*_*i*_ are held constant, exp *z*_1_ and exp *z*_2_ will be concave functions of *a*_*i*_; such diminishing-return relationships are commonly observed in genotype-phenotype-fitness maps [55, 56].

Mutations in the *cis*-element can affect *a*_1,*i*_ and/or *a*_2,*i*_ but not *b*_*i*_; *b*_*i*_ can only be affected by mutations in downstream genes, which are not being modeled in this study. The effects of mutations in the *i*-th *cis*-element on log-transformed binding affinities (ln *a*_1,*i*_ and ln *a*_2,*i*_) follow a bivariate normal distribution *N* (**0, m**_*i*_), where

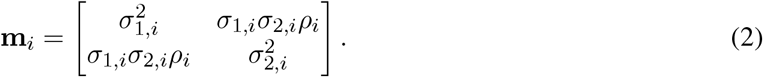

The diagonal elements of **m**_*i*_ are the variances of mutations’ effects on ln *a*_1,*i*_ and ln *a*_2,*i*_, respectively, and *ρ*_*i*_ is the correlation between the effects of mutations on ln *a*_1,*i*_ and ln *a*_2,*i*_. The effect of a mutation can be represented as a vector 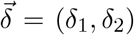 sampled from *N* (**0, m**_*i*_). After the mutation, *a*_1,*i*_ and *a*_2,*i*_ become *a*_1,*i*_ exp (*δ*_1_) and *a*_2,*i*_ exp (*δ*_2_), respectively. The mutational variance-covariance matrix for **z** = [*z*_1_ *z*_2_] ^⊤^is a sum of mutational input across the *cis*-elements [57, 58]:

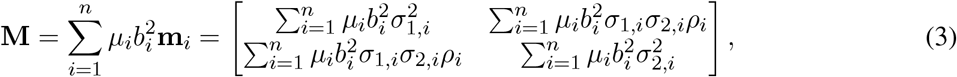

where *µ*_*i*_ is the mutation rate of the *i*-th *cis*-element per haploid genome.

A *cis*-element may be non-pleiotropic (regulating effector gene expression in only one body part) or pleiotropic (regulating effector gene expression in more than one body part) [59–61]. In this study, three classes of *cis*-elements are considered: the first includes *cis*-elements that are deployed exclusively in *Z*_1_, referred to as *Z*_1_-specific *cis*-elements; similarly, those deployed exclusively in *Z*_2_ are referred to as *Z*_2_-specific *cis*-elements; the last includes *cis*-elements that are deployed in both body parts, referred to as pleiotropic *cis*-elements. The numbers of *cis*-elements in these three classes are denoted as *n*_1_, *n*_2_, and *n*_*p*_, respectively. A *Z*_1_-specific *cis*-element has *σ*_2,*i*_ = 0, and similarly, a *Z*_2_-specific *cis*-element has *σ*_1,*i*_ = 0. For a pleiotropic *cis*-element, *σ*_1,*i*_ and *σ*_2,*i*_ are of the same order of magnitude.

In this study, for simplicity, we let all *cis*-elements contribute the same amount of mutational variance per trait: there is *σ*_1,*i*_ = *σ* and *σ*_2,*i*_ = *σ* for *Z*_1_-specific and *Z*_2_-specific *cis*-elements, respectively, *σ*_1,*i*_ = *σ*_2,*i*_ = *σ* for pleiotropic *cis*-elements, and *µ*_*i*_ = *µ* and *b*_*i*_ = *b* for all *cis*-elements. We also assumed *ρ*_*i*_ = *ρ* for all pleiotropic *cis*-elements, which means *ρ* reflects only the difference in binding preference of the two *trans*-factors. We only considered values of *ρ >* 0, which is interpreted as that *R*_1_ and *R*_2_ are paralogous and have at least somewhat similar binding preference, as paralogous TFs are known to specify identities of repeated body parts (e.g., the *Hox* genes [2, 27, 62, 63]). Mutational variances for *z*_1_ and *z*_2_ are thus V_1_ = *µ*(*n*_1_ + *n*_*p*_)*b*^2^*σ*^2^ and V_2_ = *µ*(*n*_2_ + *n*_*p*_)*b*^2^*σ*^2^, respectively, and the mutational covariance is Cov(*z*_1_, *z*_2_) = *µn*_*p*_*b*^2^*σ*^2^*ρ*. Taken together, the **M**-matrix can be written as

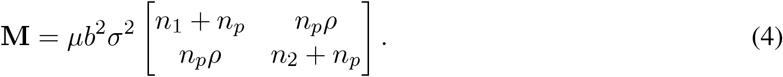

In a diploid population, the genetic covariance matrix **G** is approximately **G** = 2*N*_*e*_**M** under neutrality and at mutation-drift balance [33, 57, 64–66]. Under the simplified numerical assumption made for Eq. 4 (to be followed for the rest of this study, unless specified otherwise), there is

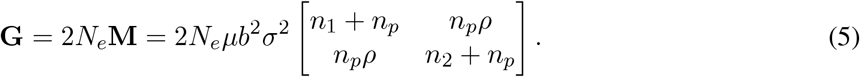

Our simulations were performed under the simple numerical assumption of Eq. (4) with *b* = 0.1, *n* = *n*_1_ + *n*_*p*_ = *n*_2_ + *n*_*p*_ = 50, *σ* = 1, and ancestrally *a*_1,1_ = *a*_1,2_ = *a*_2,1_ = *a*_2,2_ = 1 (such that *z*_1_ = *z*_2_ = 0), unless specified otherwise. Values of *n*_*p*_ that were considered include 0, 10, 20, 30, 40, and 50. We considered *ρ* = 0.9 (a minor difference in binding preference between *trans*-factors) and *ρ* = 0.5 (a moderate difference in binding preference between *trans*-factors).

### Mutation accumulation simulations

To illustrate the effect of developmental constraints on mutational variances and covariances, we simulated 200 mutation accumulation (MA) lines for each combination of *n*_*p*_ and *ρ*, under the simple numerical assumption of Eq. (4). The number of mutations acquired by each MA line was set to 5*n*_1_ + 5*n*_2_ + 5*n*_*p*_, with each *cis*-element having equal probability of being hit by a mutation. The expected phenotype variance among MA lines at the end of the simulation is thus equal to 5*nb*^2^*σ*^2^ = 2.5 for both *z*_1_ and *z*_2_. At the end of the simulations, for each combination of *n*_*p*_ and *ρ*, we calculated Pearson’s correlation coefficient between *z*_1_ and *z*_2_ across MA lines, and compared it to the theoretical expectation (given by 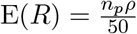).

### Evolutionary dynamics under selection

Fitness is modeled as a multivariate Gaussian function of the bivariate phenotype **z** = [*z*_1_ *z*_2_] ^⊤^, and the strength of selection along different phenotypic dimensions was characterized by a covariance matrix **S**. Fitness, denoted as *ω*, is calculated as [33, 64, 67]

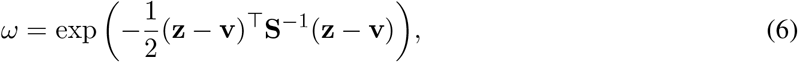

where the column vector **v** is the optimal phenotype.

When genetic variance is high and selection is relatively weak (the diagonal elements of **G** are much smaller than the corresponding elements of **S**), the evolutionary dynamics of the population mean phenotype can be approximated by an Ornstein–Uhlenbeck process (OU) model [33]. Its instantaneous evolutionary rate at time *t* is thus given by

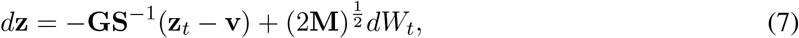

where *dW*_*t*_ denotes a unit Brownian motion process. Neutral evolutionary change over time *T* accordingly follows a bivariate normal distribution *N* (**0**, 2**M***T*). Each diagonal element of **GS**^*−*1^ is the attraction parameter of the corresponding phenotypic trait and should be between 0 and 1. If values of **G** and **S** result in attraction parameters beyond the range, the OU approximation will no longer hold.

### Simulating the evolution of the population mean phenotype

We simulated 20 replicate lineages (populations) for each combination of parameter values and examined the mean phenotype across populations over time. Simulation time was divided into time steps. Mutation rate per *cis*-element per haploid genome per time step was set to 5 *×*10^*−*5^; this setting can be interpreted as a per-generation mutation rate of 10^*−*7^ with each time step corresponding to 500 generations. The mutational variance of each trait per haploid genome per time step is thus *V*_*M*_ = 2.5*×*10^*−*5^. Diagonal elements of the **S**-matrix were *S*_1,1_ = *S*_2,2_ = 100 in all scenarios. We simulated evolution with *N*_*e*_ = 10^5^ and **G** = 2*N*_*e*_**M**, which means the **G**-matrix per time step has diagonal elements *G*_1,1_ = *G*_2,2_ = 5. The diagonal elements of **GS**^*−*1^ are then equal to 0.2, which is appropriate for the attraction parameter of an OU process, allowing the OU approximation of Eq. (7). The population mean phenotype at the end of the *t*-th time step was obtained by

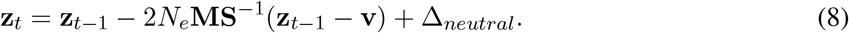

The **M**-matrix here is the per time step mutational variance-covariance matrix and Δ_*neutral*_ is sampled from *N*(**0**, 2**M**). All simulations started with an ancestral state **z**_*a*_ = [0 0] ^⊤^ (numerical assumption of Eq. (4) with all elements of **a** equal to 1 and *b* = 0.1).

We considered three scenarios of selection involving directional selection on *z*_1_. In all three scenarios, there is *v*_1_ = ln 10 (10-fold difference in non-normalized trait value from the ancestral state). In the first scenario, *z*_2_ is under stabilizing selection (*v*_2_ = 0) and there is no correlational selection (*S*_1,2_ = *S*_2,1_ = 0). In the second scenario, there is selection on the proportion between *z*_1_ and *z*_2_ and the optimal allometric relationship is *v*_1_ = *v*_2_, such that *v*_2_ = 10 and *S*_1,2_ = *S*_2,1_ = 90. In the last scenario, there is a functional tradeoff between *z*_1_ and *z*_2_ and the optimal allometric relationship is *v*_1_ + *v*_2_ = 0, such that *v*_2_ = *−* ln 10 and *S*_1,2_ = *S*_2,1_ = *−*90.

### Population genetic simulation

Population genetic simulations were conducted in SLiM [68]. We simulated diploid Wright-Fisher populations of relatively small size (*N* = 1000) but under strong directional selection (*v*_1_ = ln 100, *S*_1,1_ = *S*_2,2_ = 1). Three scenarios of selection were considered: in the first scenario, *v*_2_ = 0 and *S*_1,2_ = *S*_2,1_ = 0; in the second scenario, where selection on proportion is present, *v*_2_ = ln 100 and *S*_1,2_ = *S*_2,1_ = 0.9; in the last scenario, where a tradeoff between *z*_1_ and *z*_2_ is present, *v*_2_ = *−*ln 100 and *S*_1,2_ = *S*_2,1_ = *−*0.9. Mutation rate was set as 10^*−*7^ per *cis*-element per haploid genome. Recombination rate was 0.5 (free recombination), which is interpreted as indicating the *cis*-elements are sparsely distributed along the chromosome. All pleiotropic *cis*-elements had *ρ* = 0.9. For each combination of regime of selection and *n*_*p*_, we simulated 10 populations for 10*N* = 10^4^ generations, calculated the mean phenotype across populations 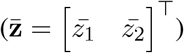over time and the corresponding fitness.

### Fluctuating selection

Evolution under fluctuating selection was simulated under an OU framework with the optimum shifting periodically. Population mean phenotype at the end of the *t*-th time step, denoted **z**_*t*_, is obtained in a similar way as Eq. (8):

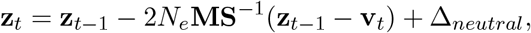

where **v**_*t*_ is the optimum for the *t*-th time step, and Δ_*neutral*_ is sampled from *N* (**0**, 2**M**). The **M** and **S**-matrices remain constant over time. At the end of the *t*-th time step, fitness corresponding to the population mean phenotype was calculated for each lineage as

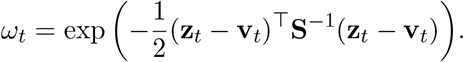

Simulations with fluctuating selection were conducted with *N*_*e*_ = 10^5^ and *ρ* = 0.9. For each regime of fluctuating selection, we simulated 100 replicate lineages for 200 time steps. Movements of **v** in different lineages were independent. To quantify the average adaptive performance over time of lineages with the same *n*_*p*_, we calculated, at the end of each time step, the mean fitness of all lineages as 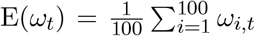, where *ω*_*i,t*_ is the fitness value corresponding to the *i*-th population’s mean phenotype at the end of the *t*-th time step. For each lineage, we calculated the geometric mean fitness over time as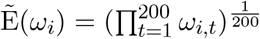 which represents the lineage’s overall adaptive performance over time.

Two models of adaptive peak movement, referred to as the Brownian motion (BM) model and the white noise (WN) model respectively, were considered.

### Brownian motion (BM)

Under this model, the movement of **v** over time is described by a BM process, and is expected to produce BM-like dynamics of phenotypic evolution as the population tracks the movement of the adaptive peak [34]. In our simulations, the movement of **v** took place at the beginning of each time step, at which time a vector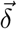 would be sampled from a bivariate normal distribution *N* (**0**, Σ_*e*_) and the optimum for the *t*-th time step would be 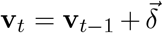. The matrix 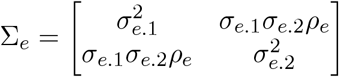describes the pattern of movement of the optimum, with the diagonal elements capturing the rate of movement and off-diagonal elements capturing internal selection that constrains the movement [6].

We considered three scenarios under this type of fluctuating selection. In the first scenario, where *z*_1_ and *z*_2_ fluctuate independently and there is no correlational selection, there is *ρ*_*e*_ = 0. In the other two scenarios, correlational selection is present, and the optimum generally moves along a selective line of least resistance (SLLR) where stabilizing selection is weakest [30, 31]. In one of these scenarios, there is selection on proportion (*S*_1,2_ = *S*_2,1_ = 90) and there is *ρ*_*e*_ = 0.9. In the other scenario, there is a tradeoff between *z*_1_ and *z*_2_ (*S*_1,2_ = *S*_2,1_ = *−* 90) and *ρ*_*e*_ = *−* 0.9. In this study, we had *v*_1_ and *v*_2_ move at the same rate and *σ*_*e*.1_ = *σ*_*e*.2_ = *σ*_*e*_. Values of *σ*_*e*_ used for simulations include 1, 2, and 4.

### White noise (WN)

Under this model, **v** is resampled from a pre-specified multivariate normal distribution *N* (**0**, Σ_*e*_) every *τ* time steps, such that **v**_*t*+*τ*_ is uncorrelated with **v**_*t*_. For our simulations, we had *τ* = 20 and the first sampling of **v** take place at the beginning of the first time step. We considered the same three scenarios of correlational selection as described above: *S*_1,2_ = *S*_2,1_ = 0 and *ρ*_*e*_ = 0, *S*_1,2_ = *S*_2,1_ = 90 and *ρ*_*e*_ = 0.9, and *S*_1,2_ = *S*_2,1_ = *−* 90 and *ρ*_*e*_ = *−* 0.9. We also had *σ*_*e*.1_ = *σ*_*e*.2_ = *σ*_*e*_, and values of *σ*_*e*_ used for simulations include 10 and 20.

All simulations and analyses of simulation results were done in R [69].

## Acknowledgment

We thank Günter Wagner, Joanna Wolfe, Joshua Schraiber, and Mark Kim for feedback on earlier versions of this manuscript. D.J. and L.S. are supported by Okinawa Institute of Science and Technology Graduate University.

## Data accessibility

Code and data files used in this study are available at https://github.com/DoubleAnonReview/evolution_repeated_structure.

## Authors’ contributions

D.J.: conceptualization, formal analysis, methodology, visualization, writing– original draft, writing–review and editing. M.P.: conceptualization, writing–review and editing. L.S.: conceptualization, writing–review and editing, Funding acquisition.

## Supplementary Materials

**Figure S1:**
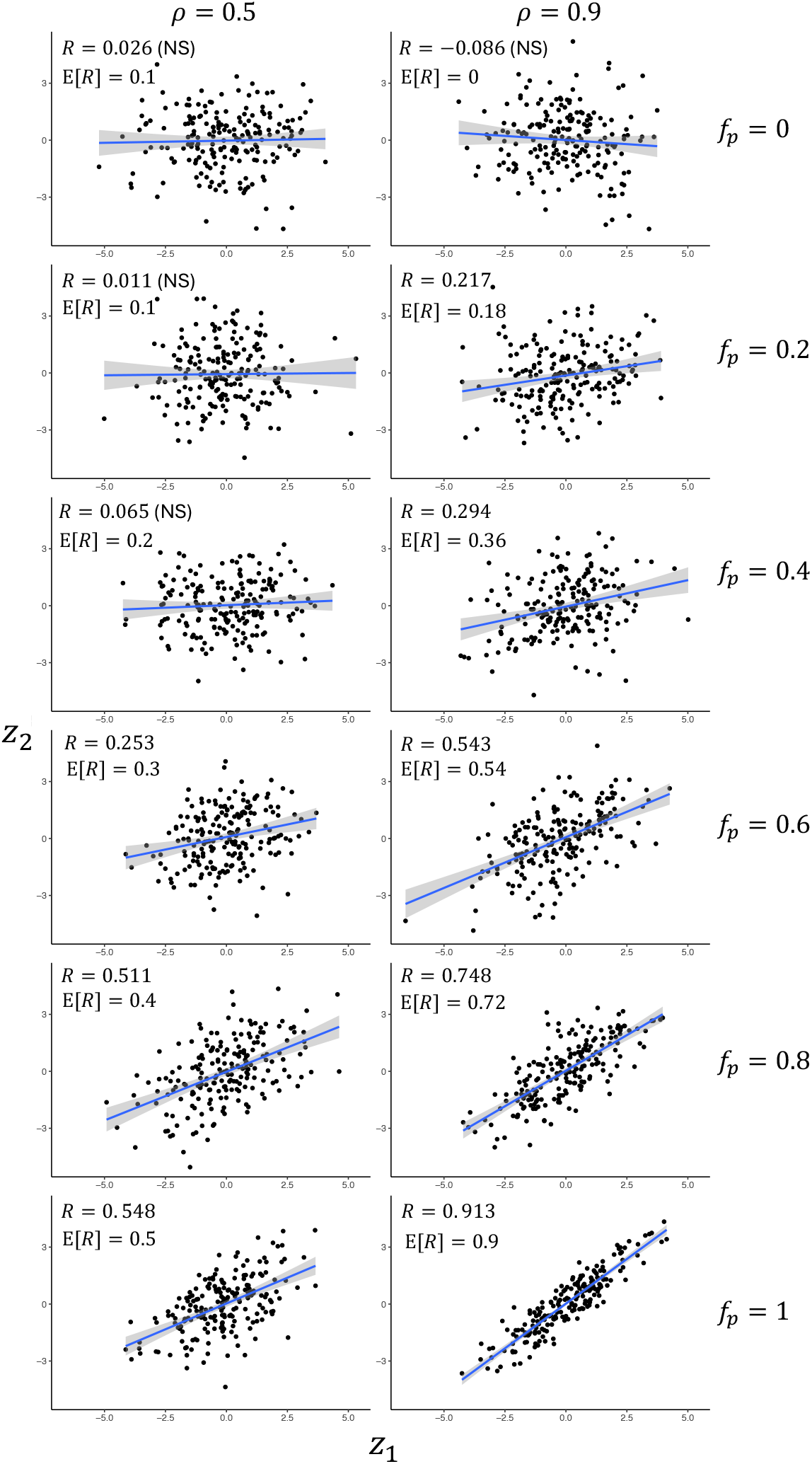
Phenotypic variation among simulated mutation accumulation (MA) lines. The number of mutations acquired in each MA line is five times the total number of *cis*-elements. Each data point corresponds to an MA line. Pearson’s correlation coefficient between *z*_1_ and *z*_2_ (*R*) and the corresponding theoretical expectations (E(*R*)) are shown in each panel, and "NS" means the observed correlation is not significant (*P >* 0.05).

**Figure S2:**
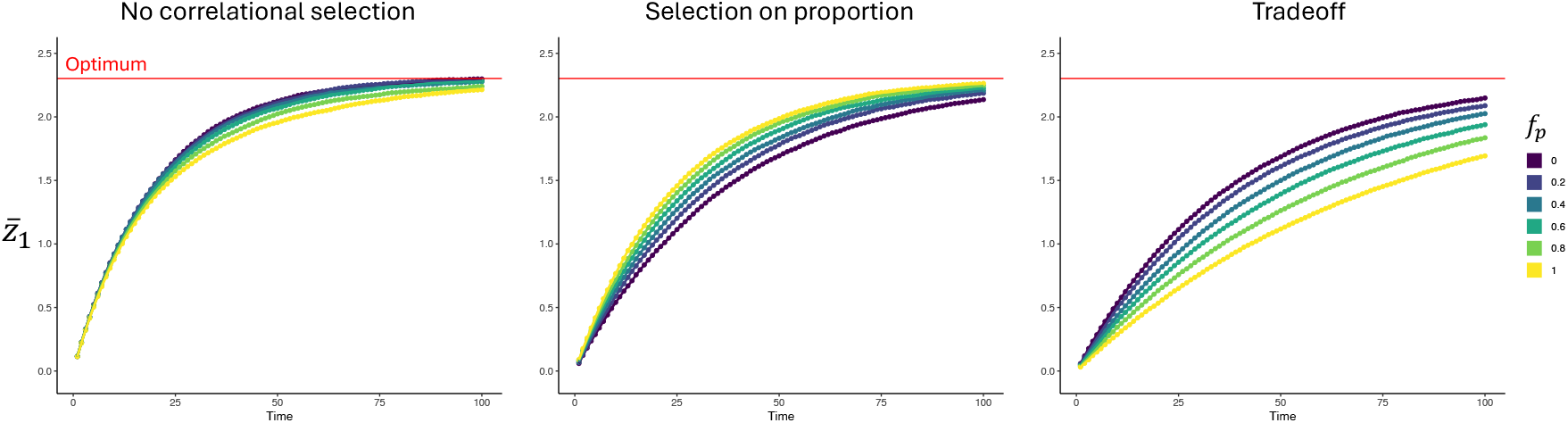
Response of the phenotype to directional selection when TFs expressed in two body parts have moderately similar binding preferences (correlation coefficient between pleiotropic mutation’s effect on *z*_1_ and *z*_2_ is *ρ* = 0.5; see Methods). The left, middle, and right panels correspond to the regimes of selection in Fig.2A.

**Figure S3:**
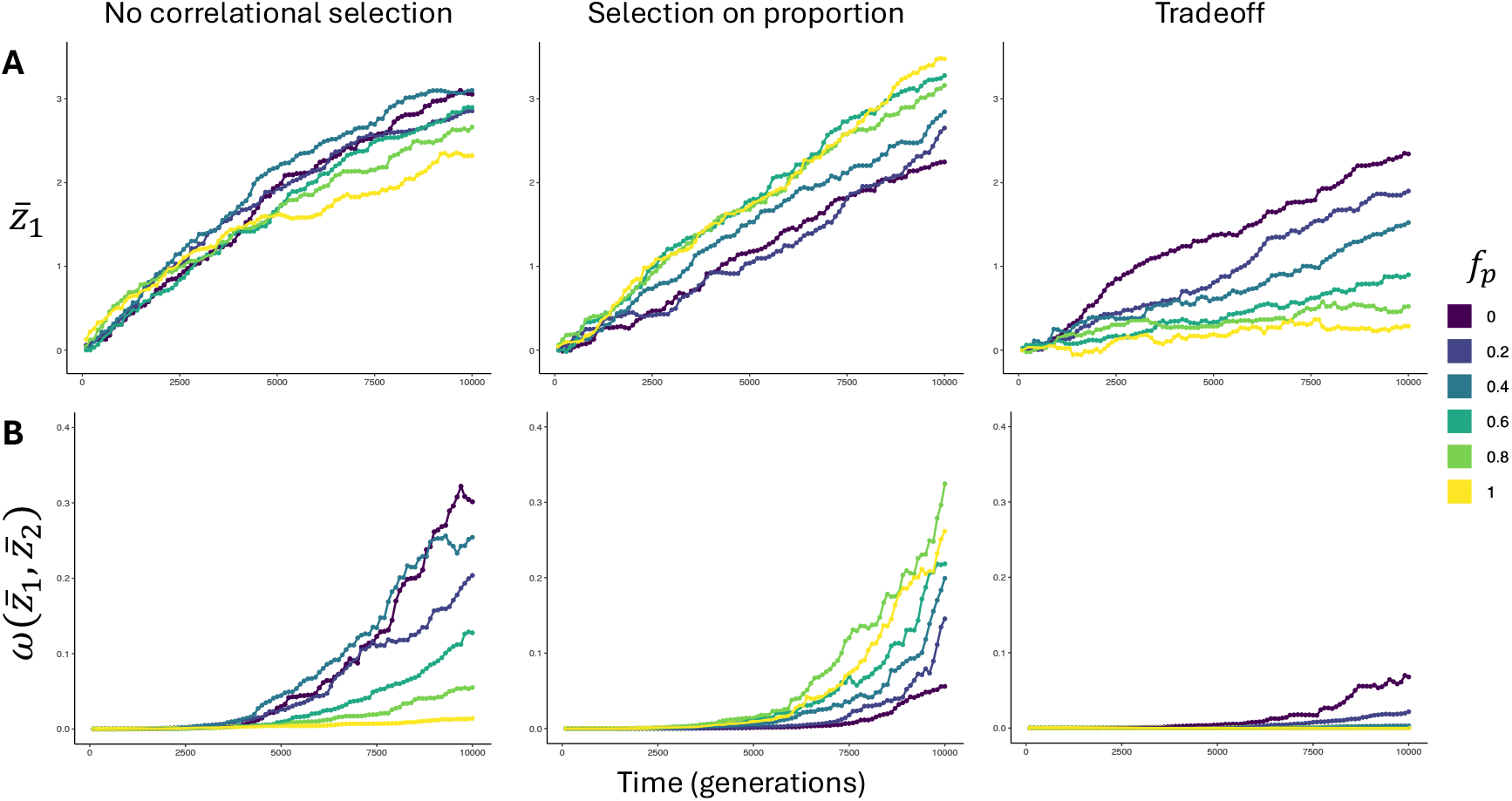
Response of the phenotype to directional selection in population genetic simulations. Left, middle, and right columns correspond to the scenarios of selection in the left, middle, and right columns in Fig.2A, respectively, except that selection is stronger (narrower fitness function and greater distance between ancestral and optimal phenotypes; see Methods). (A) Mean of z1 across populations 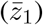 over time. (B) Fitness corresponding to the mean phenotype across populations 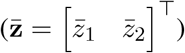 over time. The population size was *N* = 1000 for all simulations.

**Figure S4:**
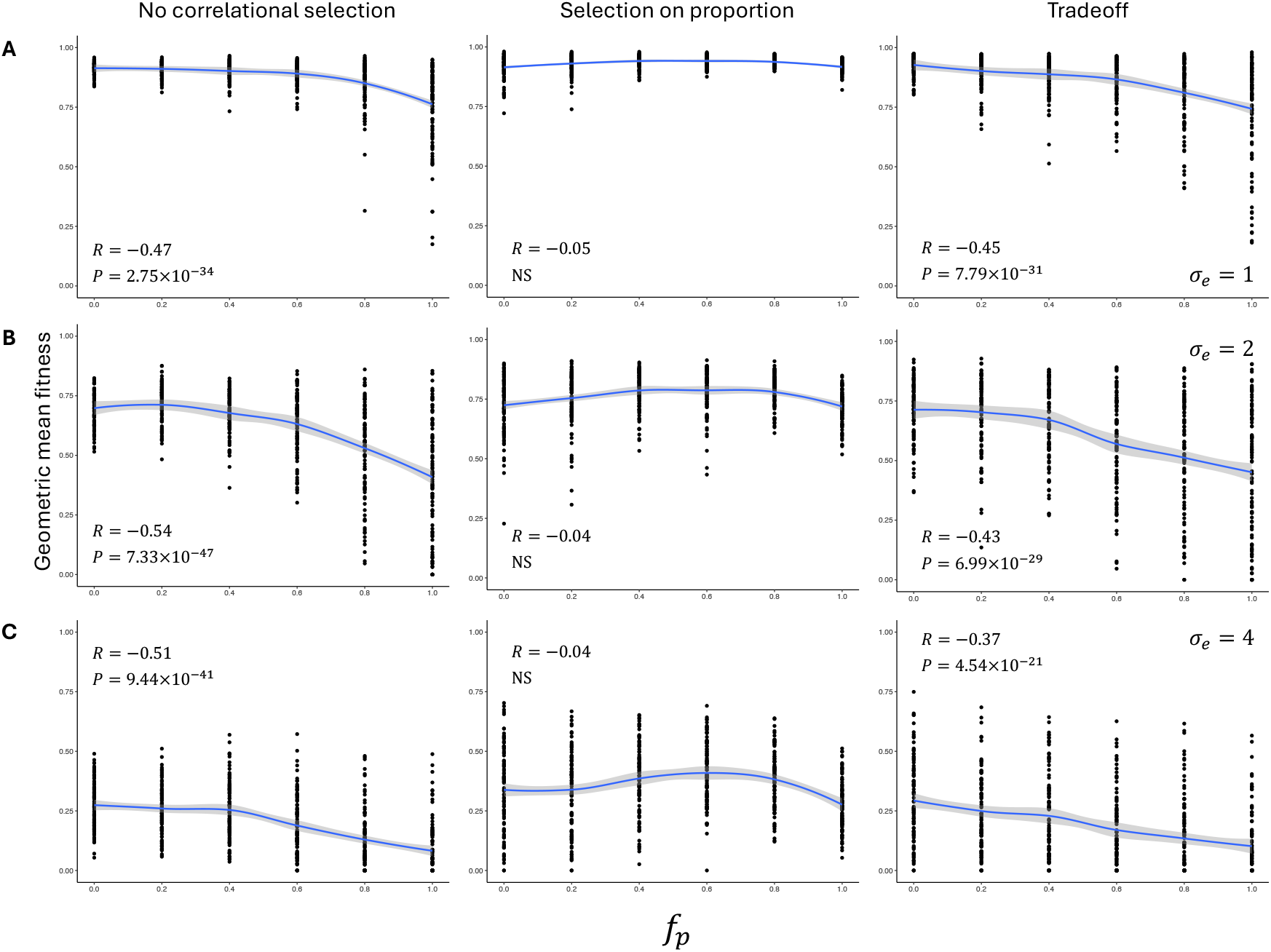
Geometric mean fitness of lineages that evolved under fluctuating selection, plotted against *f*_*p*_. The adaptive peak moved according to BM models, and the left, middle, and right columns correspond to modes of peak movement in the left, middle, and right panels of Fig. 3A, respectively. Each data point corresponds to a simulated lineage. Curve in each panel is a LOESS curve and the gray area around the curve is the 95% confidence interval. Pearson’s correlation coefficient between the geometric mean fitness and *f*_*p*_ along with the *P*-value (or "NS" if *P* > 0.05) is is noted in each panel. (A) The adaptive peak moves at a relatively slow rate (*σ*_*e*_ = 1). (B) The adaptive peak moves at an intermediate rate (*σ*_*e*_ = 2). (C) The adaptive peak moves at a relatively fast rate (*σ*_*e*_ = 4).

**Figure S5:**
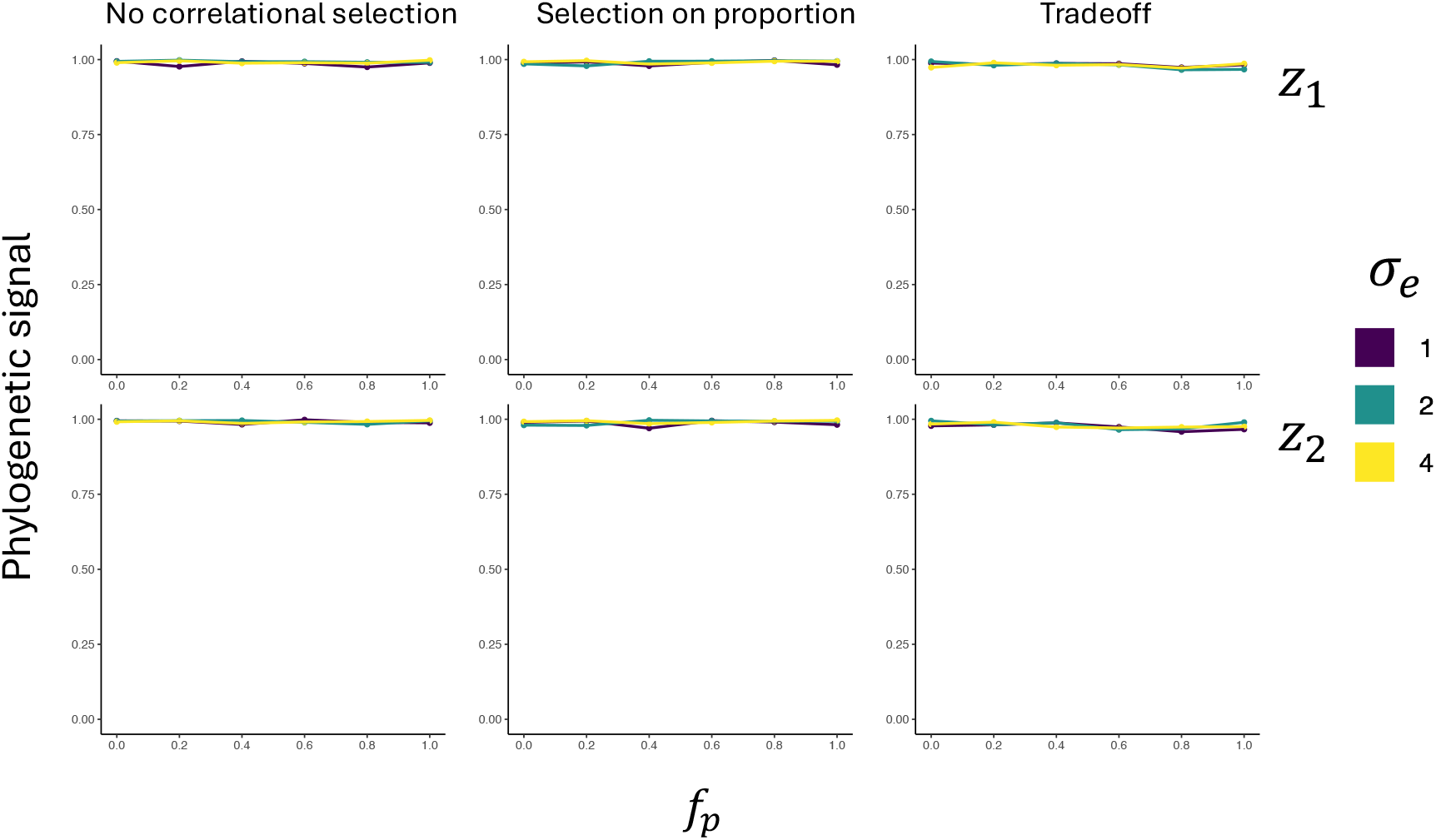
Phylogenetic signal of *z*_1_ (top row) and *z*_2_ (bottom row), quantified as Pearson’s correlation coefficient between time and phenotypic variance among populations. The adaptive peak moved according to BM models, and the left, middle, and right columns correspond to modes of peak movement in the left, middle, and right panels of Fig. 3A, respectively.

**Figure S6:**
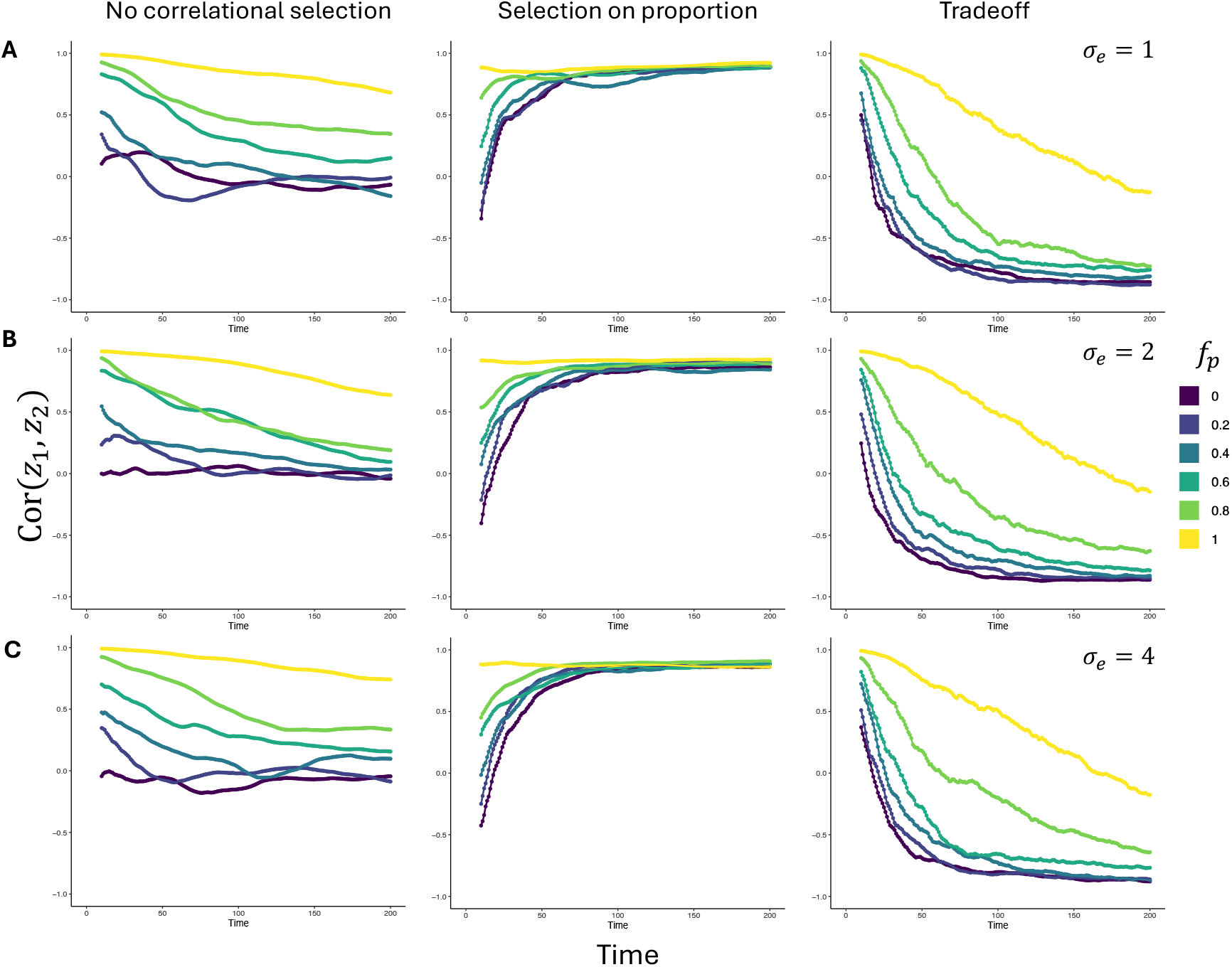
Correlation between *z*_1_ and *z*_2_ across lineages over time while the adaptive peak moved following BM models. The left, middle, and right columns correspond to modes of peak movement in the left, middle, and right panels of Fig. 3A, respectively. (A) The adaptive peak moves at a relatively slow rate (*σ*_*e*_ = 1). (B) The adaptive peak moves at an intermediate rate (*σ*_*e*_ = 2). (C) The adaptive peak moves at a relatively fast rate (*σ*_*e*_ = 4).

**Figure S7:**
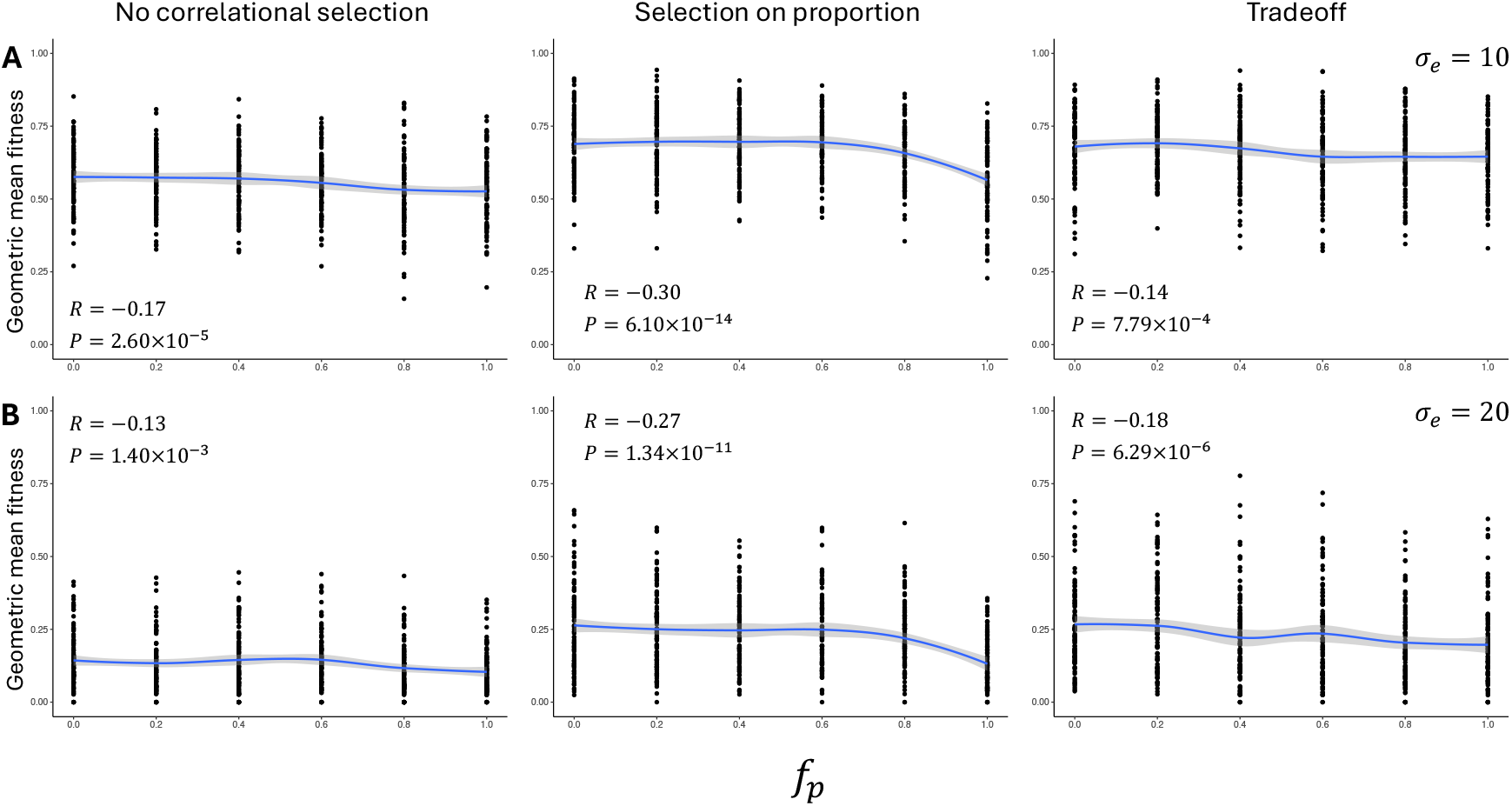
Geometric mean fitness of lineages that evolved under fluctuating selection, plotted against *f*_*p*_. The adaptive peak moved according to memory-less WN models, and the left, middle, and right columns correspond to modes of peak movement in the left, middle, and right panels of Fig. 4A, respectively. Curve in each panel is a LOESS curve and the gray area around the curve is the 95% confidence interval. Pearson’s correlation coefficient between the geometric mean fitness and *f*_*p*_ along with the P-value (or "NS" if *P* > 0.05) is is noted in each panel. (A) The optimum is resampled from a relatively narrow distribution (standard deviation *σ*_*e*_ = 10). (B) The optimum is sampled from a relatively wide distribution (standard deviation *σ*_*e*_ = 20).

**Figure S8:**
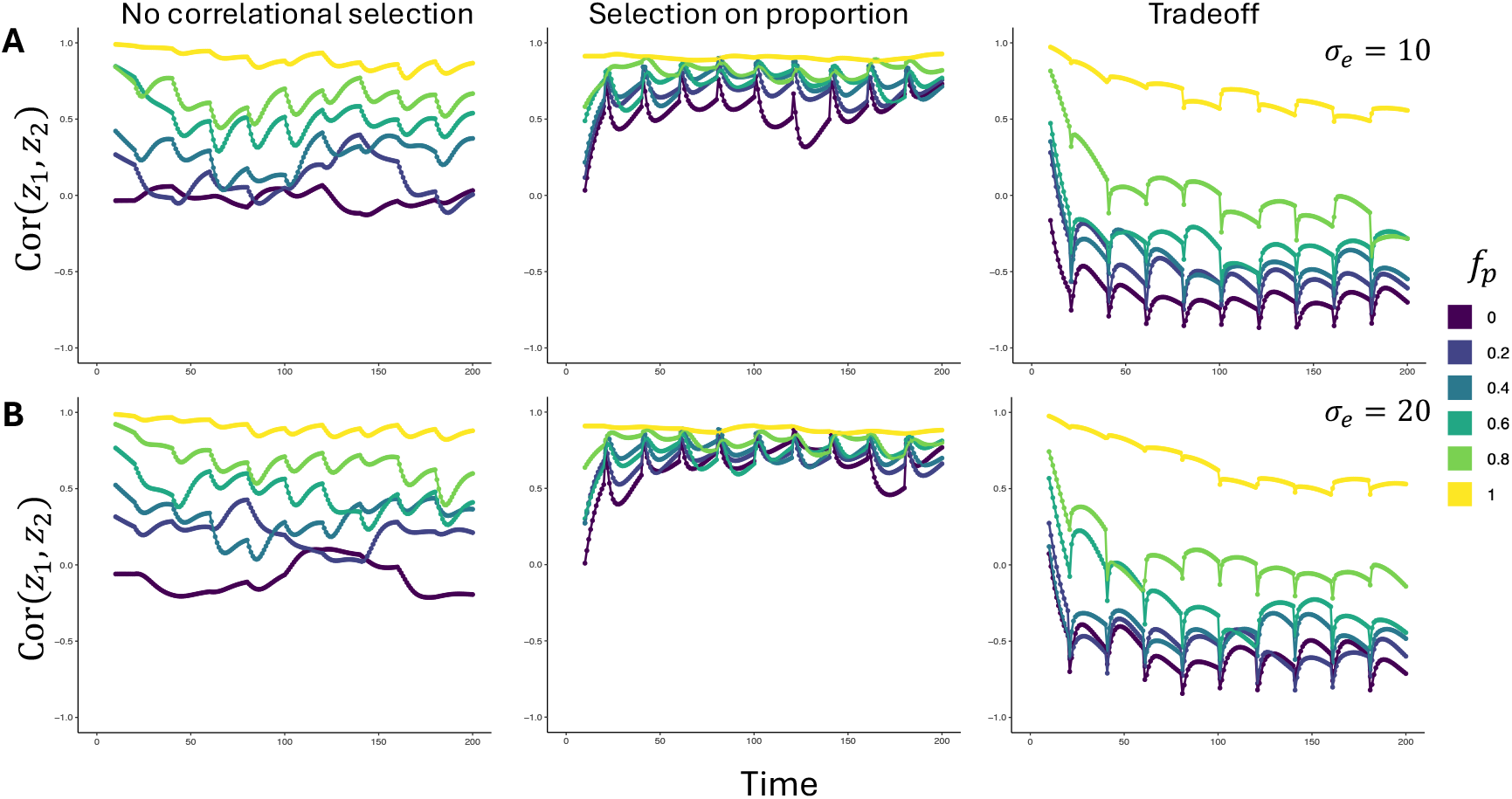
Correlation between *z*_1_ and *z*_2_ across lineages over time while the adaptive peak moved according to memory-less WN models. The left, middle, and right columns correspond to modes of peak movement in the left, middle, and right panels of Fig. 4A, respectively. (A) The optimum is resampled from a relatively narrow distribution (standard deviation *σ*_e_ = 10). (B) The optimum is sampled from a relatively wide distribution (standard deviation *σ*_e_ = 20).

